# Dynamics, Electrostatics, and Thermodynamics of Base Pairing at the *LTR-III* Quadruplex:Duplex Junction

**DOI:** 10.1101/2024.01.17.576154

**Authors:** Haley M. Michel, Justin A. Lemkul

**Affiliations:** Department of Biochemistry, Virginia Tech, Blacksburg, Virginia, 24061, United States; Center for Drug Discovery, Virginia Tech, Blacksburg, Virginia, 24061, United States

## Abstract

G-quadruplexes (GQs) play key regulatory roles within the human genome and have also been identified to play similar roles in other eukaryotes, bacteria, archaea, and viruses. Human immunodeficiency virus 1 (HIV-1), the etiological agent of acquired immunodeficiency syndrome (AIDS), can form two GQs in its long terminal repeat (LTR) promoter region, each of which act to regulate viral gene expression in opposing manners. The major LTR GQ, called *LTR-III*, is a distinct hybrid GQ containing a 12-nucleotide duplex loop attached to the quadruplex motif. The resulting quadruplex:duplex junction (QDJ) has been hypothesized to serve as a selective drug targeting site. To better understand the dynamics of this QDJ, we performed conventional and enhanced-sampling molecular dynamics simulations using the Drude-2017 force field. We observed unbiased and reversible formation of additional base pairs in the QDJ, between Ade4:Thy14 and Gua3:Thy14. Both base pairs were electrostatically favored but geometric constraints within the junction may drive the formation of, and preference for, the Ade4:Thy14 base pair. Finally, we demonstrated that the base pairs are separated only by small energy barriers that may enable transitions between both base-paired states. Together, these simulations provide new insights into the dynamics, electrostatics, and thermodynamics of the *LTR-III* QDJ.

**SIGNIFICANCE:** Here, we characterize the quadruplex:duplex junction of the HIV-1 *LTR-III* G-quadruplex. We find that two additional base pairs can form in the junction and are driven by electrostatic, thermodynamic, and geometric factors. G-quadruplexes containing such junctions are rather recent discoveries, and it has been proposed that these junctions can act as selective targets for drugs. These results further identify distinct chemical and electrostatic characteristics that can be used to guide drug design studies.

## INTRODUCTION

G-quadruplexes (GQs) comprise a class of noncanonical nucleic acid structures that form in guanine-rich sequences in both DNA and RNA. GQ-forming sequences are non-randomly distributed throughout the genomes of eukaryotes, bacteria, archaea, and viruses and are particularly enriched in regions that regulate gene expression and cell and viral survival. As such, GQs can play distinct functional roles in the etiology of human diseases and indicate that GQs can potentially act as selective therapeutic targets for a variety of neurodegenerative disorders (1), cancers (2), and bacterial (3) and viral infections (4). GQs that form in viral sequences are understudied compared to those found in the human genome but nearly all human-infecting viruses are enriched with GQ-forming sequences that are conserved within viral classes (5). The lentivirus genus, which is responsible for many severe and fatal immunological diseases and includes human immunodeficiency virus 1 (HIV-1), relies on a conserved and tightly regulated viral promoter within the 5’-long terminal repeat (LTR) that contains GQ-forming sequences (6, 7). These GQs have been shown to be key control mechanisms for viral replication and may provide insight into the viral regulation that can be used to create more effective anti-viral therapeutics (8–10).

In HIV-1, the etiological agent of acquired immunodeficiency syndrome, viral replication is regulated through multiple GQs found in both the viral and proviral genomes (8, 9, 11, 12). Two RNA GQs, called *U3-III* and *U3-IV*, regulate reverse transcription through the U3 domain of the viral RNA genome (10, 13). The structures of these RNA GQs have yet to be experimentally determined, but the analogous DNA GQ structures found in the U3 region of the proviral 5’-LTR have been resolved by NMR (14, 15). These mutually exclusive GQs, denoted *LTR-III* and *LTR-IV*, inversely regulate LTR promoter activity and expression of all HIV-1 genes (8). *LTR-III* is the predominant structure and its formation induces promoter silencing, whereas *LTR-IV* formation leads to promoter enhancement (14, 15). While this behavior ultimately implicates *LTR-III* as the primary nucleic acid drug target, it is also believed that *LTR-IV* stabilization has potential to over-activate the replicative mechanism and cause premature cell death prior to virion production, thus impacting viral latency in the host. The interplay between these two distinct GQs is thought to be modulated by protein binding and highlights a finely tuned viral regulatory mechanism that can be altered through selective targeting of one GQ structure over the other (13, 16).

Thus far, therapeutic targeting of GQs has been limited due to a lack of compounds that selectively target GQs of interest. The main structural feature of GQs is the guanine tetrad core, which is composed of stacked layers of four guanines in a planar organization that are stabilized by Hoogsteen hydrogen bonds. These tetrads, present in all GQ structures, serve as the main interaction site for current GQ-targeting compounds. However, the regions flanking the tetrad guanines can form distinct and highly dynamic loop regions that may aid in the selective targeting of individual GQ structures. In regards to the LTR GQs, *LTR-III* has a unique 12-nucleotide duplex loop coaxial to the quadruplex motif (Figure 1). Between these motifs lies a quadruplex:duplex junction (QDJ), a feature that has been suggested to serve as a selective drug target in GQ structures (17–19). Until recently, the idea of QDJs was merely hypothesized and most studies revolved around synthetically designed quadruplex:duplex hybrid constructs to test their ability to be selectively targeted (20–24). Several QDJs have been hypothesized to form in the promoters of human oncogenes like *BCL-2, RAB3D*, and *RAB12* (17) while few others such as *PIM1* (25) have been structurally determined. The base composition and length of these duplex loops can vary and ultimately influence the stability of the duplex (26) and the folding kinetics of the entire GQ structure (27). The majority of QDJs contain base pairs stacked on top of a guanine tetrad and the orientation of these bases can partially block the electronegative GQ core, a common ligand binding site, thus modulating the chemical properties of this junction and ultimately its accessibility to small molecules (22). *LTR-III* contains a distinct QDJ that is believed to be dynamic and can form different base pairs (15). Targeting the junction of *LTR-III* has been of interest to several groups and has shown promising results in regards to selective targeting, but the dynamics of the QDJ may complicate targeting studies. Therefore, the potential geometric and biophysical properties of the junction may provide a key insights that can be translated for effectively targeting this structure.

**Figure 1:**
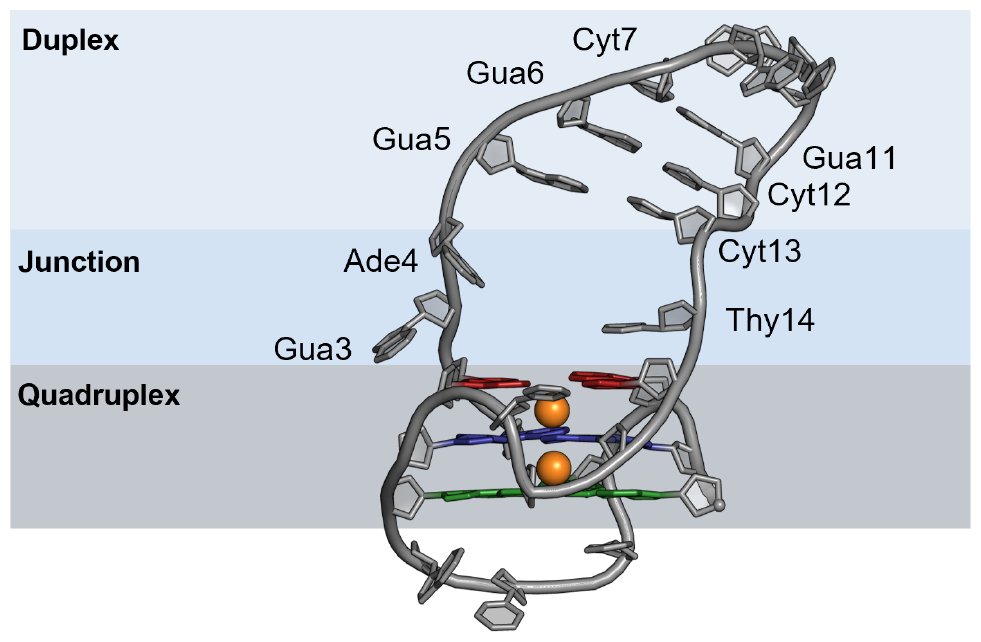
Structure of the *LTR-III* GQ. The structure was obtained from model 1 of the NMR ensemble deposited in PDB entry 6H1K. The structure is characterized by a 12-nucleotide duplex loop (light blue) containing three canonical Gua:Cyt base pairs (labeled). The junction between the quadruplex and duplex motif is open in the NMR model and contains three bases, Gua3, Ade4, Thy14 (labeled). The prototypical GQ tetrad core is shown with three tetrads (red, blue, and green) ligating two K^+^ ions (orange spheres).

Here, we build upon our previous studies (28–31) employing the Drude-2017 polarizable force field (FF) (32, 33) to study the dynamic QDJ of *LTR-III*. We use conventional and Gaussian-accelerated molecular dynamics simulations to explore how variations in base hydrogen bonding patterns within the junction impact the electrostatic, thermodynamic, and geometric properties of this potential drug binding site.

## METHODS

### System Construction

The starting structure for the *LTR-III* GQ was taken from the first model of the NMR ensemble in PDB entry 6H1K (15). Two K^+^ ions were added to the GQ core in bipyramidal antiprismatic coordination using the same method described previously (28). The *LTR-III* GQ was centered in a cubic box with a minimum box-solute distance of 10 Å. These boxes were filled with CHARMM-modified TIP3P water molecules and a total salt concentration of ∼ 150 mM, including neutralizing K^+^ counterions. Once constructed, the solvated system was relaxed via energy minimization in CHARMM (34) through 500 steps of steepest descent minimization and 500 steps of adopted-basis Newton-Raphson minimization. Equilibration was performed for 1 ns under an *NPT* ensemble in NAMD (35). All non-hydrogen and the core K^+^ were restrained for the duration of equilibration while water and bulk KCl were free. The system was equilibrated at 298 K, maintaining the temperature with the Langevin thermostat method (36) with a friction coefficient of 5 ps^-1^. The pressure was kept constant at 1 atm by the Langevin piston method (37). Periodic boundary conditions were applied in all spatial dimensions. The short-range van der Waals forces were smoothly switched from 10 to 12 Å. The particle mesh Ewald method (38, 39) was used to calculate electrostatic interactions with a real-space cutoff of 12 Å, and 1-Å Fourier grid spacing. All bonds to hydrogen were constrained using the SHAKE algorithm (40) while the water molecules were held rigid through the SETTLE algorithm (41). The integration timestep for the nonpolarizable sytems is set to 2 fs.

### Conventional MD Simulations with the Drude-2017 FF

The final coordinates of the nonpolarizable equilibration were used to prepare Drude systems. Drude oscillators and lone pairs were added to all heavy atoms in the system with the CHARMM program. TIP3P water molecules were converted to the polarizable SWM4-NDP model (42) and the Drude-2017 FF (32, 33) was applied to the GQ and ions. Drude oscillators were relaxed through energy minimization with all real atoms held fixed, allowing the induced dipoles to relax. During equilibration, the Drude oscillators were coupled to a low-temperature relative thermostat at 1 K with a friction coefficient of 20 ps^-1^. The nonbonded treatment was also the same as the nonpolarizable equilibration except the van der Waals potential was switched from 10 to 12 Å rather than the van der Waals forces. Heavy atoms were again restrained and a “hard wall” constraint was applied to allow a maximum Drude-atom bond length of 0.2 Å to avoid polarization catastrophe. Equilibration of the Drude systems was carried out for 1 ns with a time step of 1 fs. Unrestrained, production simulations were then performed in OpenMM (43) for 1 *μ*s per replicate, resulting in a total simulation time of 4 *μ*s..

### NOE Distance Validation

Hydrogen-hydrogen distances were extracted from Nuclear Overhauser Effect (NOE) spectroscopy data deposited in the NMR restraint files associated with PDB 6H1K. We then separated out all internucleotide NOEs for use in validation. Intranucleotide NOEs were omitted as they describe sensitive conformational changes rather than overall agreement in the structural ensemble, as we have done previously (31). Thymine atoms labeled C7 were also excluded, as the hydrogens in these NOEs are not uniquely defined. In total, we evaluated 239 H-H NOE distance ranges for *LTR-III* and compared those to the same atom distances within the simulations.

### Electric Field Analysis

Electric field vectors were calculated with TUPÃ (44), using “atom” mode with the center-of-mass of each base of interest (Gua3, Ade4, and Thy14) acting as the probe point. The environment consisted of the entire GQ structure and any water molecules and KCl ions within 20 Å of the probe. The magnitude of the field and the vector points in the x-, y-, and z-directions were calculated and reported every 100 ps. The alignment of the electric field vector, 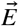, with the corresponding base dipole moment vector, 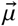, was then calculated to characterize the response of the selected base to its electrostatic environment. Electric field vectors run in the direction of the movement of positive charge while dipole moment vectors were represented in the chemical convention and point from partial positive charge to partial negative charge.

### Gaussian-Accelerated MD Simulations

Dual-boost Gaussian-accelerated MD (GaMD) was applied to the unequilibrated Drude systems with each simulation consisting of 2 ns conventional MD, 50 ns GaMD equilibration, and 600 ns GaMD production simulation in NAMD (45). The threshold energy, *E*, was set as the lower bound, where *E=V*_*max*_, or the maximum potential and dihedral energies. The default parameter values were used for GaMD simulations unless otherwise stated. The window for statistical averaging of the potential energies and the frequency of boost recalculation was set to 400 ps. Convergence of GaMD equilibration was assessed by monitoring *k*_*0*_ and the threshold energy, *E*, over time. Production runs were started using the atomic velocities from the last step of equilibration. The GaMD simulations were energetically reweighted to recover the potential mean of force (PMF) profiles using cumulant expansion to the second order within the PyReweighting toolkit (46). The angle formed between the centers-of-mass of the Ade4, Gua3, and Thy14 nucleobases and duplex eRMSD were used as collective variables to reweight the simulation data and calculate the free energy surface of *LTR-III*. PMF convergence was verified via block reweighting every 100 ns.

## RESULTS AND DISCUSSION

### Dynamic Base Pairing at the QDJ Occludes the GQ Core

*LTR-III* adopts a hybrid quadruplex:duplex topology characterized by a 12-nucleotide duplex loop containing three canonical base pairs and three bases at its QDJ that are unpaired in the NMR ensemble (Figure 1). During the conventional MD simulations, the duplex region adopted several conformations with different relative positioning over the GQ core and with differing degrees of twisting. To identify the dominant states in this conformational ensemble, we performed RMSD-based clustering in CHARMM. The algorithm uses a self-organizing neural net to cluster data points based on a maximum radius from the cluster center (47–49). Clusters were generated by pooling all replicates and separating frames based on a maximum radius of 3 Å. The least-occupied cluster (Cluster 7, Figure S1), representing 1% of total simulation time, was characterized by duplex conformations that resembled the original NMR model, in which the QDJ is open and the duplex loop is flat. The majority of the other clusters contained twisted duplex loops that resemble B-DNA due to the three canonical base pairs present in the loop. Unlike the NMR models, the most-occupied clusters exhibited closed junctions, such that the solvent-accessible cavity at the QDJ is occluded. The most-occupied cluster (Cluster 1, Figure S1) contains an additional base pair between Ade4 and Thy14 within the QDJ, whereas the second most-occupied cluster lack this base pair, but the duplex loop shifts down to close off the junction (Cluster 2, Figure S1).

Analysis of the individual trajectories revealed that clusters containing the closed junction without any base pairing arose from replicates 1 and 4; the Ade4:Thy14 base-paired states only occurred in replicates 2 and 3. Additionally, a noncanonical Gua3:Thy14 base pair formed in the junction when Thy14 was not paired with Ade4. The Gua3:Thy14 pair was the first to form in both replicates and acted as a scaffold for Thy14 to then shift upward to pair with Ade4. Ade4:Thy14 was the more prevalent base pair, appearing in about 38% of the total 4-*μ*s simulation time, whereas Gua3:Thy14 was only present for 10% (Figure 2B). Interestingly, the Gua3:Thy14 pair adopted two different states, with Gua3 in either *syn* or *anti* orientations. In replicate 2, Gua3 was *syn* and formed two hydrogen bonds, involving Gua3(H1):Thy14(O4) and Gua3(O6):Thy14(H3), as shown in Figure 2A. In replicate 3, Gua3 was in the canonical *anti* orientation, resulting in a bifurcated hydrogen bond with Gua3 H1 and H21 acting as donors with Thy14 O4 as the acceptor (Figure 2C). Both base pairs occluded the junction but were also directly above a solvent-facing tetrad, a common site for drug targeting and ion binding (28, 50). These base pairs were found to limit ion access to these sites, suggesting that they may also impact small-molecule binding at this location (Figure S2).

**Figure 2:**
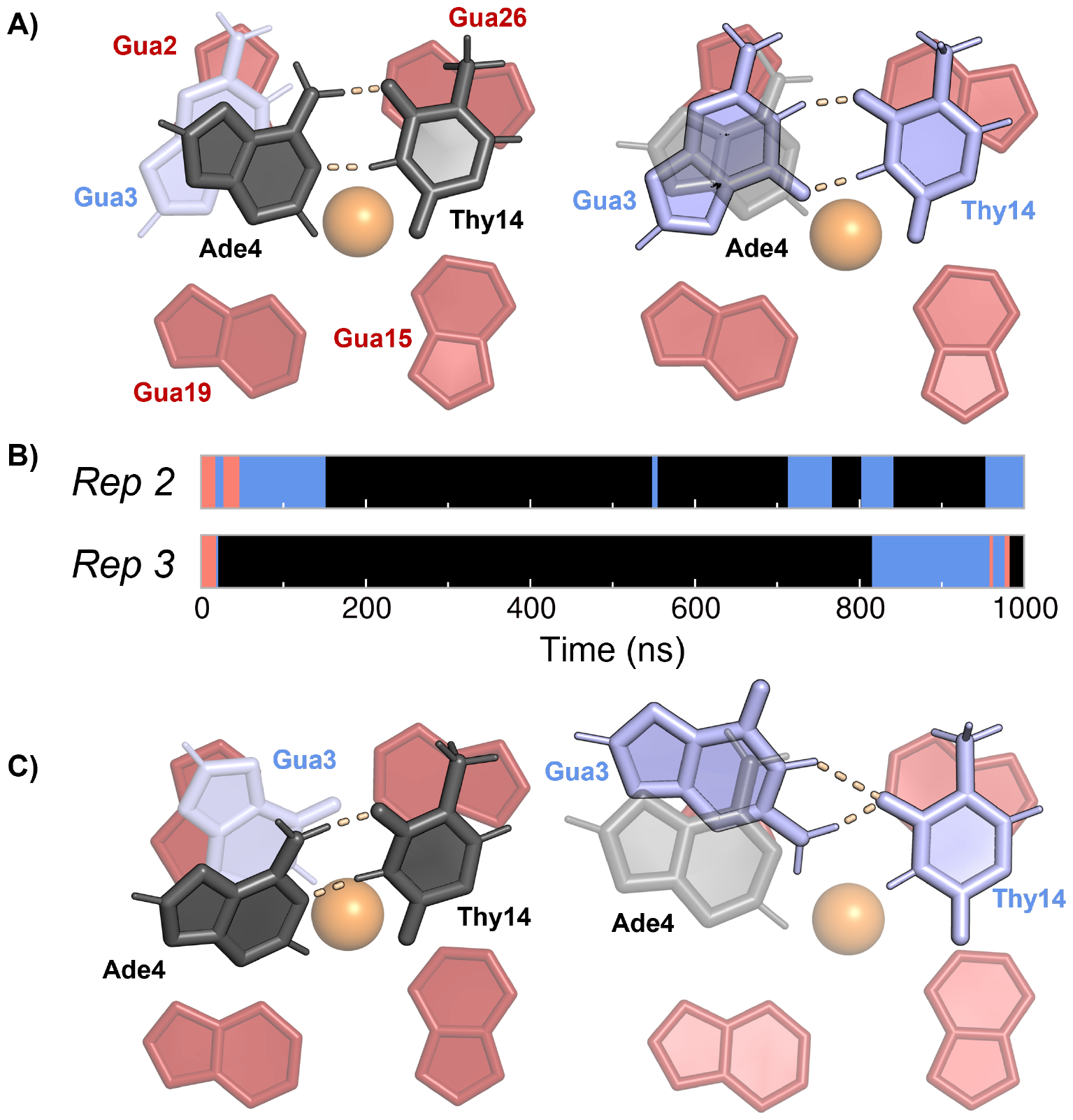
Hydrogen-bonding patterns in the QDJ. Both Ade4:Thy14 (black) and Gua3:Thy14 (blue) form directly above tetrad 1 (red) and the GQ core, which contains K^+^ ions (orange). Base orientation and hydrogen bonds (dotted lines) are shown for both pairs in (A) replicate 2 and (C) replicate 3. (B) Time series indicating formation of each base pair and the duration that they remained formed in replicates 2 and 3.

The lack of consistent base pairing across all four replicates suggests kinetic trapping occurring in our simulations. That is, collapse of the duplex onto the quadruplex motif may have occurred faster than the inward rotation of Gua3, Ade4, and Thy14. To test this possibility, two additional simulations were executed at 310 K to compare the kinetics at 298 K to those at the higher temperature. One simulation was equilibrated as a new system from the starting NMR structure and the second simulation used the condensed duplex structure from Cluster 2 to represent the closed junction with no base pairing, as was observed in replicates 1 and 4. Both simulations exhibited junction reopening, allowing for formation of Ade4:Thy14 base pairs.Gua3:Thy14 base pairs were absent in the additional replicates at 310 K, presumably due to it being the less dominant base pair and these simulations quickly arriving at the preferred state.

These base pairs were previously investigated by Butovskaya et al. (15), who found that only Ade4:Thy14 was compatible with experimental NOEs and therefore likely to influence duplex stability. The QDJ of *LTR-III* was difficult to resolve from the available NMR restraints (A.T. Phan, personal communication), which further supports the possibility of dynamic base pair switching seen in our simulations. The formation of Ade4:Thy14 was also observed in simulations of *LTR-III* performed using the AMBER FF (51). Our calculations of NOE violations across all replicates showed high violation percentages for some inter-hydrogen distances involving Gua3, Ade4, and Thy14 (Figures S3 - S5). Similar values were also noted in the study using the AMBER FF, indicating that our results are not a FF-dependent issue but rather reinforce the difficulty in obtaining clear NMR data for the QDJ.

A better understanding of what drives the dynamics of these base pairs, including the impact of the chemical microenvironment at the QDJ, is required to further improve selective targeting efforts against *LTR-III*. Given the fact that hydrogen bonding, a largely electrostatic and strongly directional phenomenon, dictates the orientation of the base pairs, we investigated the extent to which electronic polarization and intrinsic electric fields influence these dynamics.

### Response to Electric Fields Drives QDJ Geometry

Biomolecules generate electrostatic environments designed for their specific biological functions, such as binding and catalysis. GQs generate electric fields that have been shown to influence K^+^ binding at the solvent-exposed faces of tetrads. These fields are impacted by both base dynamics and microenvironment (52). Hydrogen bonding is a largely electrostatic and directional phenomenon that dictates the orientations of the base pairs seen in the QDJ, therefore we investigated the extent to which electronic polarization influences junction dynamics. We calculated the electric field, 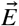, exerted by the entire GQ and nearby solvent onto Gua3, Ade4, and Thy14 using TUPÃ (44). We then determined the angle, *α*, between 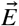 with the associated base dipole moment vector, 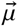, and expressed this value as the cosine of *α*. By convention, the direction of 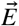 indicates the flow of positive point charges. The Drude oscillators on the heavy atoms are negatively charged, therefore we expect that the Drude oscillators would move in opposition to 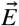. Thus, the preferred orientation of a nucleobase would be driven by an opposing alignment of 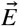 and 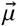, that is, *α* = 180° and *cos*(*α*) = −1.

The alignment of 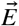 and 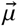 varied across replicates 2 and 3 for Gua3 and Thy14. For Gua3, the alignment of 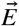 and 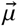 was greater in replicate 2 in the absence of base pairing at the junction compared to replicate 3, in which better alignment was achieved upon Gua3:Thy14 pairing. Thy14 showed better alignment during Ade4:Thy14 pairing in replicate 2 while differences between the alignment during Ade4:Thy14 and Gua3:Thy14 base pairing were more subtle during replicate 3 (Figure 3B). The changes in alignment preferences across replicates may be due to the different orientations of Gua3. In replicate 3, Gua3 was *anti* and may be more suitable for base pairing because it is similar to its preferred B-DNA orientation (Figure 3A). Unlike Gua3 and Thy14, Ade4 was less sensitive to base pairing in both replicates, suggesting that its base dipole moment is more resilient to the fluctuations in the surrounding electric field. We repeated the 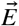 calculations without solvent contributions and found that water acts as an insulator, weakening the magnitude of the 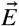 and decreasing alignment between 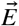 and 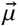. Despite the change in alignment, the trends in the calculations without solvent contributions were similar to those observed in calculations that included solvent contributions (Supporting Information Figure S6).

**Figure 3:**
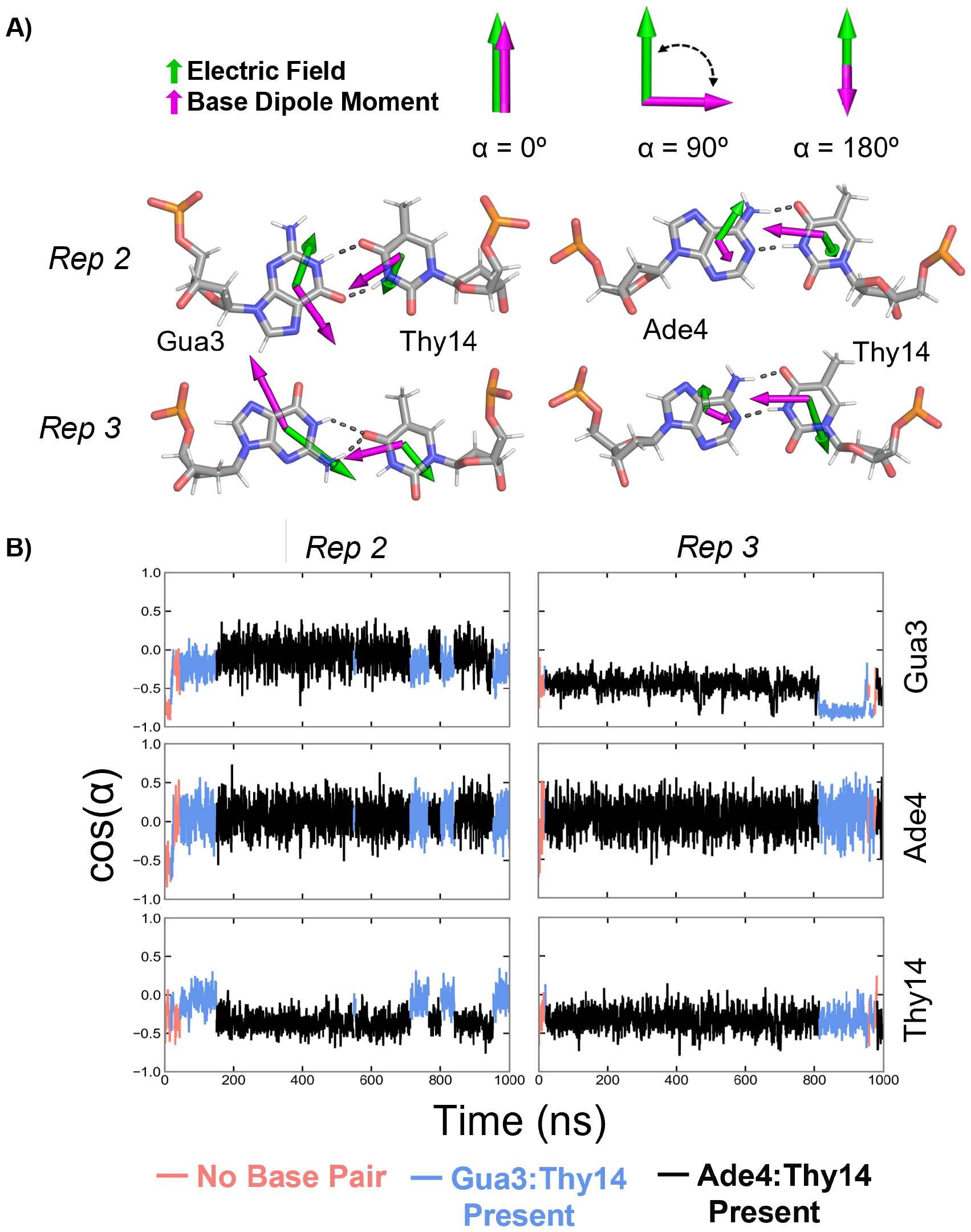
Electric field alignments for QDJ base pairs. (A) Electric field (green) and base dipole moment (pink) vectors for each base during base-pair formation. (B) Time series of the cosine of the alignment angle between the electric field and base dipole moment vectors in replicates 2 and 3. Data are plotted as running averages over 1-ns windows.

The trends in 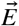 and 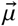 alignment indicate the preferred conformation for each individual base rather than the structure as a whole. Gua3 manifested a preference for pairing with Thy14 over the Ade4:Thy14 pair but the indifference to pairing conformations of Ade4 and the differences in Thy14 alignment across replicates 2 and 3 raise the question of why these bases do not show explicit favorability for the canonical Ade4:Thy14 pairing. Gua:Thy base pairing was previously determined to be one of the most stable mismatches, which may help explain the lack of clear preference for Ade4:Thy14 (53). Additionally, the base dipole moments of both Ade4 and Thy14 during base pairing were aligned better with each other than with 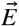. We anticipated that the 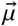 and 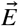 vectors would generally align, but there are other forces that act on the bases, such as bonded interactions and torsional forces that influence the geometry. Therefore, although 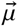 of each base may be driven to orient in a specific way due to 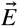, additional forces may compete with the electrostatic forces to prevent perfect alignment of 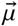 with 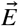.

### Base Pair Properties in the QDJ Microenvironment Resemble B-form Duplex DNA

Although the 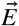 and 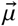 alignment indicated the preferred orientations of each individual base, we investigated the alignment of Ade4:Thy14 further, to study the specific influence of hydrogen bonding between the two bases. Total dipole moments, the summation of the permanent and induced dipoles, are influenced by more than just intramolecular electrostatic contributions that are typically targeted during parametrization. The surrounding electrostatic environment and geometric constraints placed on the constituent residues will influence the total dipole response and thus the net molecular dipole moment. The Drude FF was developed to reproduce both dipole moment and molecular polarizability computed from gas-phase QM calculations, yielding a robust model for the response to perturbing charges. A previous study with the Drude FF showed that total nucleobase dipole moments are sensitive to the surrounding microenvironment, and induced dipole contributions rationalize free energy differences between base-paired and extrahelical states (54). These differences in microenvironment may explain the orientation of the base dipole moment vectors and thus the changes in 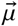 and 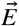 alignment for Gua3, Ade4, and Thy14 across replicates. The fact that these base pairs form at the interface between canonical duplex and guanine tetrad structural motifs suggests that the properties of the constituent bases reflect this unusual microenvironment.

To assess this possibility, we analyzed the base dipole moment, 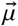 of each base and calculated the angle, *θ*, between 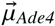 and 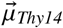 and between 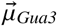 and 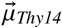 during Ade4:Thy14 and Gua3:Thy14 base pair formation and when the bases were unpaired. Only small differences in directionality and magnitude of 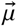 can be seen as a result of the respective change in junction conformation, which may reflect the minimal change in the surrounding microenvironment (Figure 4A). The directionality of both Ade4 and Thy14 bases in isolation have been shown using QM electrostatic potential surfaces and indicated that in a Ade:Thy base pair, 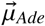 and 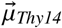 are pointed towards their respective base pairing partner (55). In this instance, *θ*_Ade-Thy_ would be close to 180°. In our simulations, 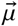 for Gua3, Ade4, and Thy14 were all found to be more aligned with their base pairing partner during Ade4:Thy14 hydrogen bonding than in any other state. Only a slight difference in *θ*_Ade4-Thy14_ arose during Ade4:Thy14 and Gua3:Thy14 pairing during replicate 3, presumably due to the differences in the orientation of Gua3 mentioned previously. Interestingly, 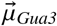 was better aligned with 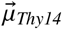 when Gua3 was not paired to Thy14. The improved alignment of 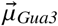 with 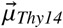 during Ade4:Thy14 pairing may indicate there are geometric constraints imposed on these bases by the DNA GQ that compete with the favorable electronic properties of Gua3 when bound to Thy14. The orientation of *θ*_Ade4-Thy14_ in our simulations was similar to that of *θ*_Ade-Thy_ in two model duplexes (Figure S9), suggesting that the Ade4:Thy14 base pairing in the QDJ resembles the stability of the same interaction in a canonical duplex.

**Figure 4:**
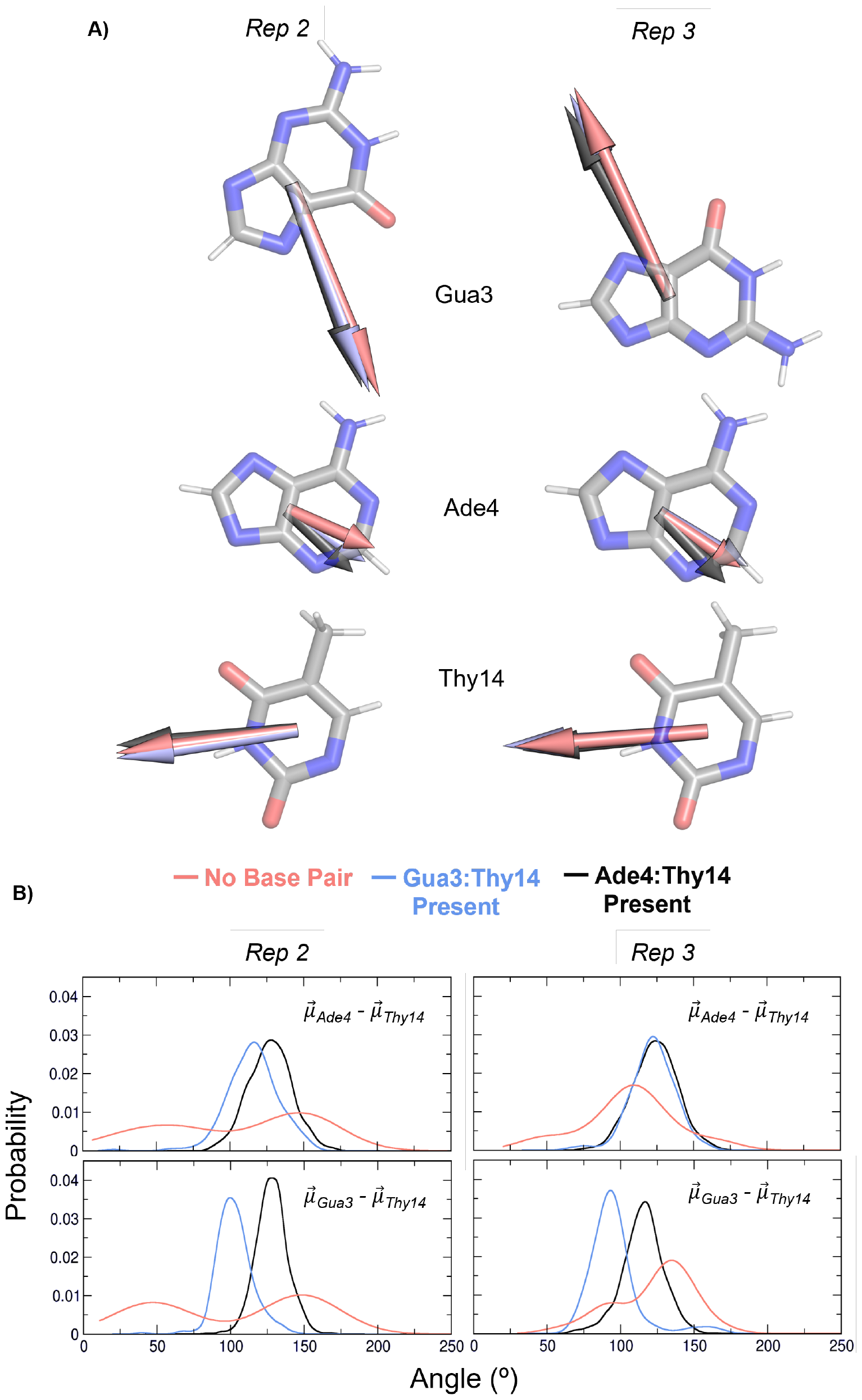
Base dipole moments of Gua3, Ade4, and Thy14. (A) Orientation of the base dipole moment vectors for states in which the base is unpaired (pink), when Gua3:Thy14 are paired (blue), and when Ade4:Thy14 are paired (black). (B) Distributions of the alignments of base dipole moment vectors between pairs of bases (Ade4 and Thy14) and (Gua3 and Thy14) when no base pairs are formed (pink), Gua3:Thy14 is formed (blue), and Ade4:Thy14 is formed (black).

Overall, these findings indicate that the entire *LTR-III* GQ structure may prefer the presence of Ade4:Thy14 in the QDJ, thus leaving Gua3 electronically frustrated as a small penalty. Base geometry has been hypothesized to work in concert with hydrogen bonding to explain the high fidelity of polymerases. In duplex DNA, the canonical base pairs have distinct geometry in regards to their C1’ distances and hydrogen bond angles, which may facilitate polymerase insertion of the correct base pair partner over other mismatched bases with similar base-pairing free energies (56, 57). These geometric constraints may apply to the duplex loop found in *LTR-III* and imply that both geometric and electrostatic properties are influential in driving the formation of different base-paired geometries in the QDJ.

### Thermodynamics of the QDJ Base Pairs

To understand the thermodynamics of the conformational transitions in *LTR-III* as a function of the different base pairing states in the QDJ, we performed GaMD simulations to construct the free energy surface associated with these states. GaMD equilibration was performed until *k*_*0*_ and *E* reached a steady value (Figure S10). We note that GaMD equilibration is unrestrained and allowed for base pair formation prior to production, therefore the resulting biasing potential was informed by the possibility of base pairing. GaMD production was then conducted for 600 ns and simulation convergence was verified performing cumulative reweighting every 100 ns (Figure S11). Similar to the hydrogen bonding patterns in conventional MD, Ade4 and Gua3 base-pairing with Thy14 was observed in the GaMD simulations. Additionally, we noted short periods of intermediate “transition states” characterized by hydrogen bonding networks among all three bases. Gua3:Thy14 was again less prevalent than the Ade4:Thy14 pair. Gua3 never sampled *syn* conformations in the GaMD simulation, but did engage in two different hydrogen bonding geometries with Thy14. These geometries were (1) bifurcated hydrogen bonds with Thy14 O4 serving as the acceptor simultaneously for both Gua3 H1 and H21 and (2) two distinct hydrogen bonds between Gua3(O6):Thy14(H3) and Gua3(H1):Thy14(O2). The ability of GaMD to provide a boost over energy barriers further revealed the formation of a tilted Gua3:Ade4:Thy14 base triple that was present for 20 ns of the simulation. During GaMD, neither Gua3 nor Ade4 rotated outward into bulk solution, but for 4 ns, neither base engaged in hydrogen bonds with Thy14. Thus, these collapsed configurations (with Gua3, Ade4, and Thy14 all directed inward along the duplex axis) are possible but disfavored compared to either base-paired state. To test whether the observed preferences of Gua3 and Ade4 were a result of kinetic trapping, we performed an additional GaMD simulation at 310 K to attempt to use simple kinetic energy to interconvert between the paired and unpaired states. This simulation confirmed that unpaired configurations, such as the condensed duplex in conventional MD, can form during equilibration but then revert to an open junction such that Ade4:Thy14 base pairs form during the production phase of GaMD.

To evaluate the thermodynamic differences between the base-paired states in our 298 K GaMD simulation, we performed 2-dimensional (2D) reweighting along collective variables defined as the eRMSD (58) of the duplex loop and the angle formed among the centers-of-mass of Ade4, Gua3, and Thy14, denoted as “AGT angle.” The resulting conformational free energy surface of *LTR-III* captured several energy basins that differentiate the Ade4:Thy14 and Gua3:Thy14 states. The majority of Gua3:Thy14 conformations were observed with AGT angles around 90° (basin A, Figure 5 and inset A2). The non-base-paired states were also found within this basin due to the planar orientation of Gua3 and Thy14 that characterize these states (Figure 5, inset A1). Ade4:Thy14 conformations were found in basins B and D, differing only in their AGT angles (Figure 5, insets B1 and D). The majority of Ade4:Thy14 base pairs were seen in basin B alongside the global minimum (basin B, marked in red in Figure 5). Also in this basin were less-occupied “transitional states” such that Thy14 is positioned between Ade4 and Gua3 to form hydrogen bonds among all three bases (Figure 5, insets B1 and B2). A final, punctate energy basin contained the distinct Gua3:Ade4:Thy14 base triple with Gua3 and Ade4 tilted on the typical base pairing axis (basin D, Figure 5). Although the global minimum corresponds to the Ade4:Thy14 base-paired states (basin B, marked in red in Figure 5), only small energy barriers, on the order of of 2-3 kcal mol^-1^, separate the base-paired states. Therefore, the GaMD results are consistent with those of the conventional MD simulations, suggesting a thermodynamic preference for Ade4:Thy14 base pairing but also indicating that the barrier to adopt Gua3:Thy14 states at 298 K can be overcome. Such a finding is important in the context of drug design, in that putative ligands that bind to the QDJ could be designed to disrupt the Gua3:Thy14 base pair. Doing so would drive the structure to form the preferred Ade4:Thy14 base pair, as the energy barrier to adopting this state can be reasonably overcome.

**Figure 5:**
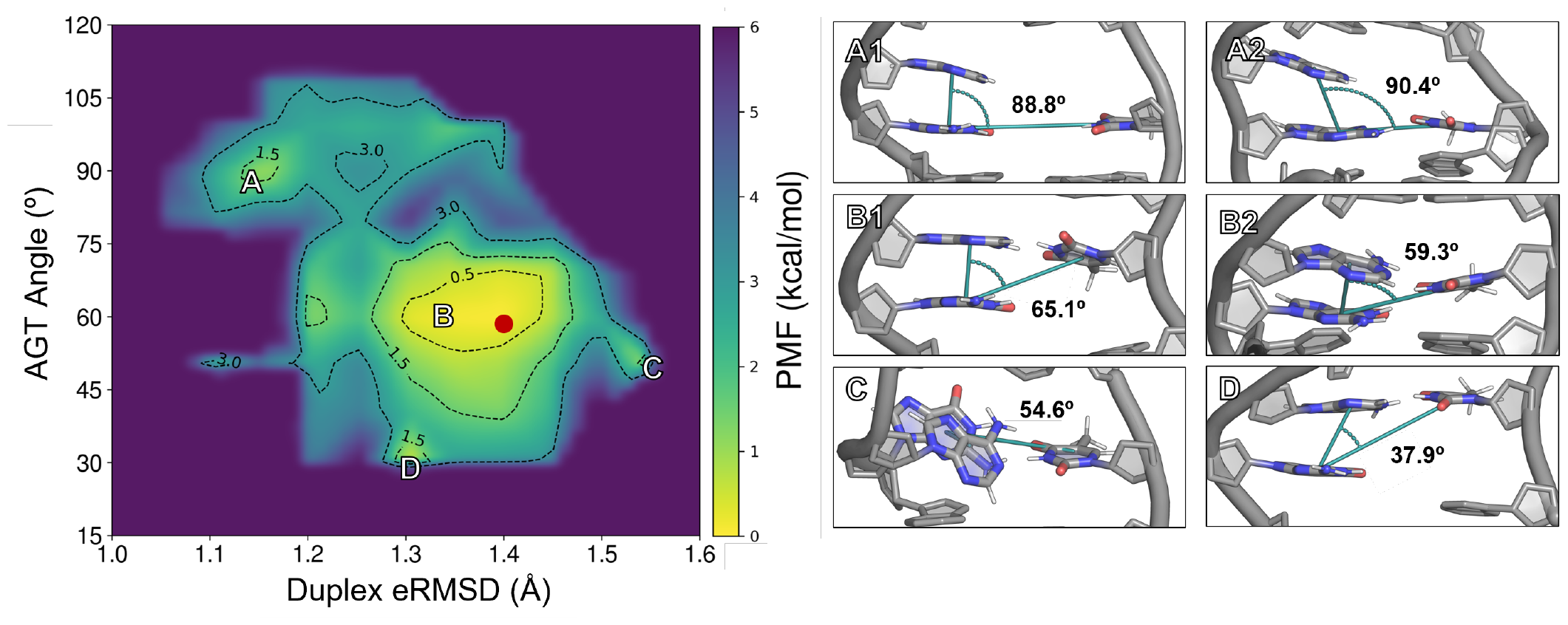
2D free energy surface associated with base pair formation. Characteristic base pairs observed in the free energy basins are indicated as (A1) no base pair, (A2) Gua3:Thy14, (B1) Ade4:Thy14, (B2) transition state, (C) base triple, and (D) Ade4:Thy14 shown in context of the QDJ. The global minimum is shown in basin B as a red dot.

## CONCLUSIONS

Here, we investigated the HIV-1 *LTR-III* GQ, with a focus on its unusual QDJ, finding that additional base pairs not modeled in the NMR ensemble form in this region of the structure. Our results support the previous hypothesis that the QDJ in *LTR-III* is highly dynamic, given that in our simulations, two different base pairs (Ade4:Thy14 and Gua3:Thy14) were observed and readily interconverted. Base-pair formation blocks the nearest tetrad in the GQ core, which leads to limited ion access, in turn suggesting that the chemical and electrostatic properties of the junction differ depending on whether the junction is solvent-accessible or occluded as a function of base-pair formation. The electric field exerted by the entire structure on the Gua3, Ade4, and Thy14 bases contributed to the favorable formation of base pairs in the QDJ, with only subtle differences between the Ade4:Thy14 and Gua3:Thy14 states. In addition to electrostatic forces, geometric constraints may play a role in dictating base-pair formation, potentially shifting preference toward Ade4:Thy14 despite the apparent electrostatic preference for Gua3 base pairing with Thy14. Only small free energy barriers separate these states, indicating the potential for facile conformational transitions within this junction. Together, this work connects the dynamics, electrostatics, and thermodynamics of distinct base-pairing conformations in the *LTR-III* GQ that provide an atomistic understanding of the forces governing plasticity at the QDJ. Moreover, the results reported here can help guide future drug design efforts by exploiting the biophysical properties and intrinsic dynamics of the junction.

## AUTHOR CONTRIBUTIONS

HMM and JAL designed the research. HMM performed and analyzed the simulations. HMM and JAL wrote the article.

## ACKNOWLEDGMENTS

The authors thank Dr. Marcelo Depolo Polêto for helpful discussions regarding data interpretation and electric fields. This work was supported by the National Institutes of Health (grant R35GM133754 to JAL) and USDA-NIFA (project VA-160092). The authors thank Virginia Tech Advanced Research Computing for computing time and resources.

## SUPPORTING INFORMATION

An online supplement to this article can be found by visiting BJ Online at http://www.biophysj.org.

