## Supporting Information for "Dynamics, Electrostatics, and Thermodynamics of Base Pairing at the *LTR-III* Quadruplex:Duplex Junction"

Manuscript submitted to **Biophysical** *Journal*

Article

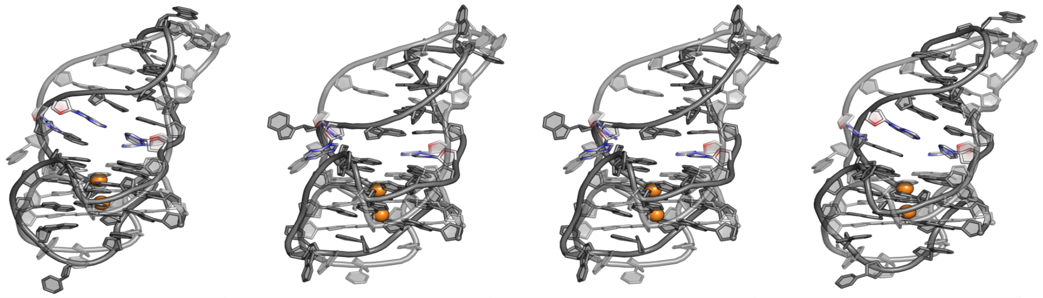

| Cluster | 1 | 2 | 3 | 4 |
|---|---|---|---|---|
| Occupancy | 21% | 20% | 19% | 16% |
| Center | 4.7 Å | 5.8 Å | 5.8 Å | 5.0 Å |

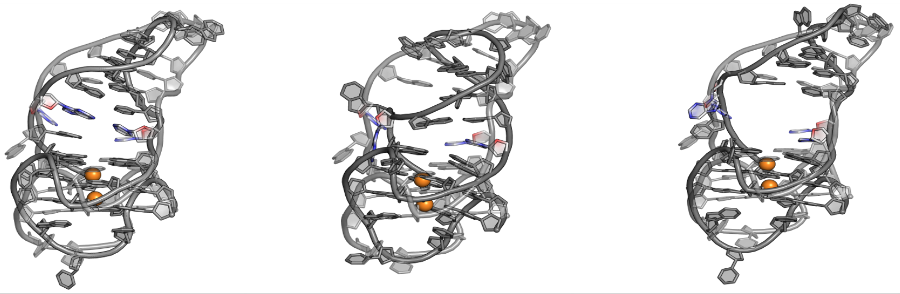

| Cluster | 5 | 6 | 7 |
|---|---|---|---|
| Occupancy | 14% | 9% | 1% |
| Center | 5.8 Å | 6.2 Å | 3.6 Å |

Figure S1: Dominant structures obtained from RMSD-based clustering of the *LTR-III* GQ. Clusters are ordered from (1) most occupied to (7) least occupied. Each cluster is overlaid on the starting structure (translucent) with the Ade4 and Thy14 base pairs colored by atom.

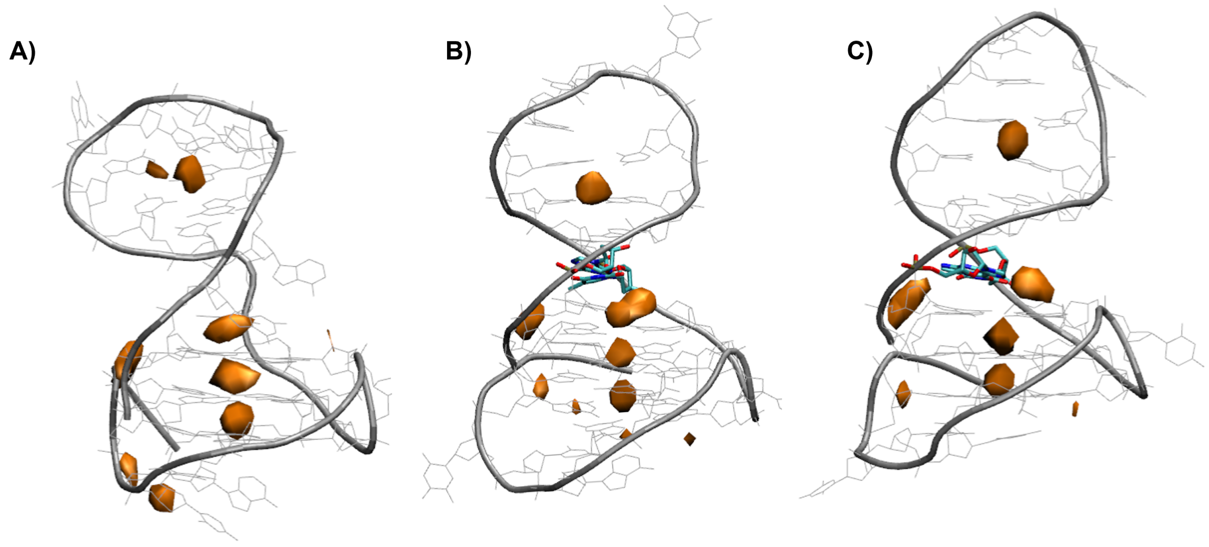


Figure S2: $K^+$ occupancy maps around the *LTR-III* GQ in the (A) no base pair, (B) Ade4:Thy14, and (C) Gua3:Thy14 states during replicate 2. The isosurface for ion sampling was set an an occupancy threshold of $\geq 1\%$. Presence of $K^+$ ions (orange) above the GQ core is altered due to the presence of base pairs (colored by atom).

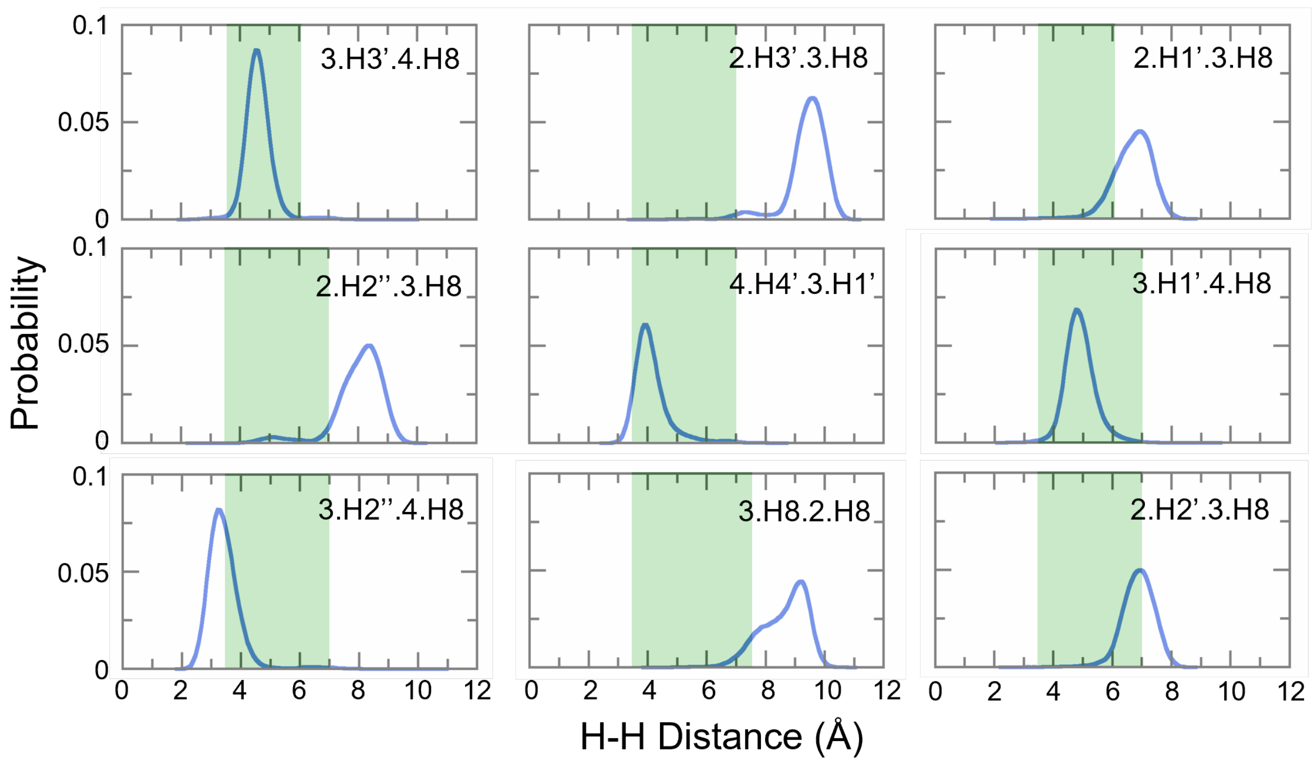


Figure S3: Gua3 internucleotide NOE distributions in replicate 2. Acceptable distance ranges (shaded green) were calculated from the raw NOE data obtained from the RCSB. Graphs are labeled with residue1.atom1.residue2.atom2 in the top right of each graph.

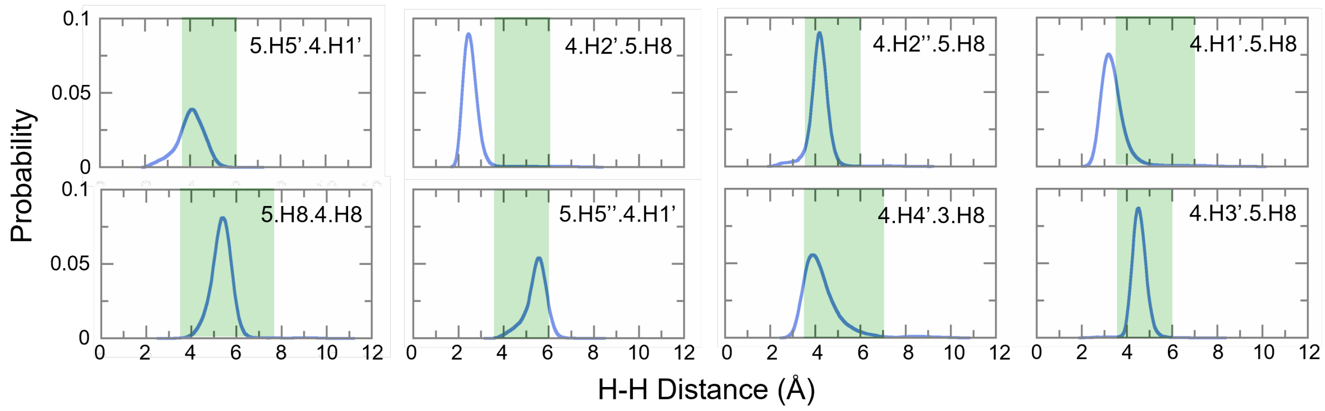


Figure S4: Ade4 internucleotide NOE distributions in replicate 2 (blue). Acceptable distance ranges (shaded green) were calculated from the raw NOE data obtained from the RCSB. Graphs are labeled with residue1.atom1.residue2.atom2 in the top right of each graph.

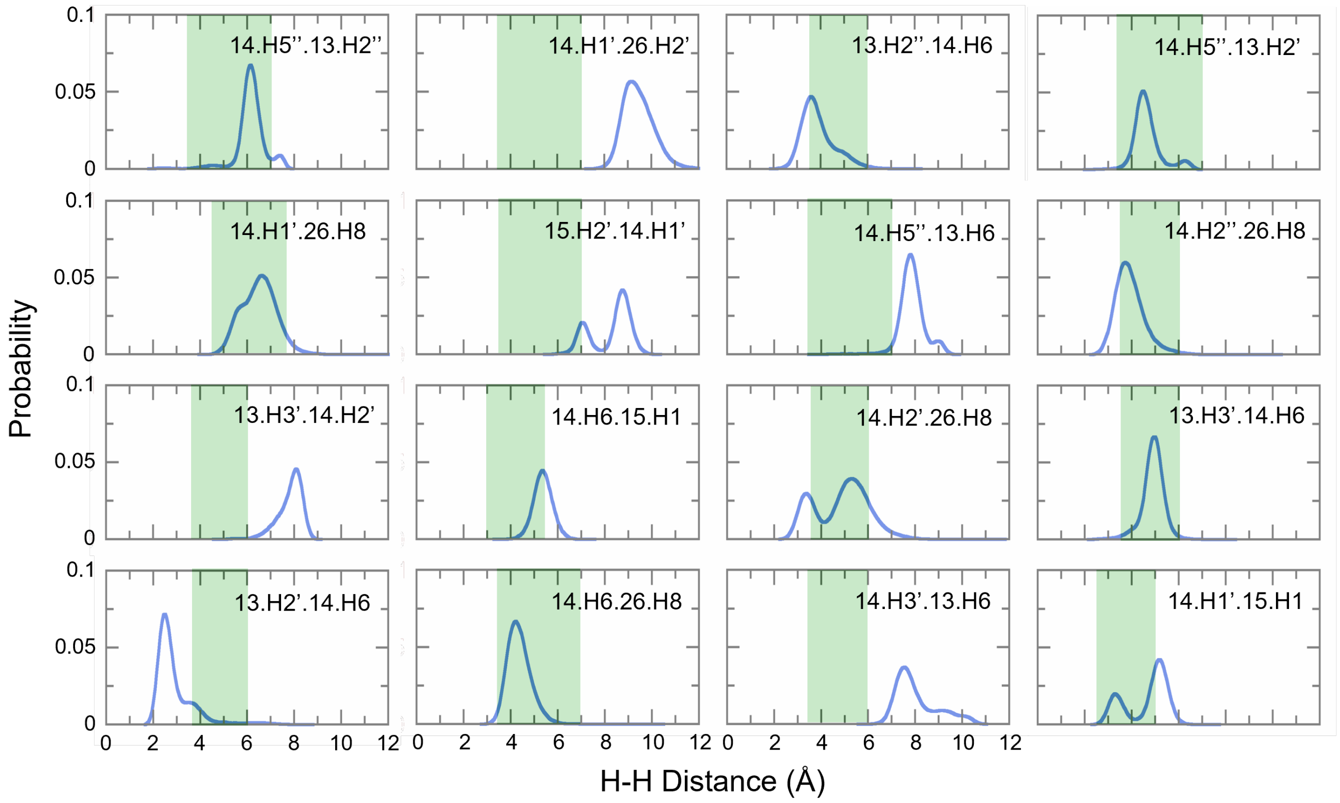


Figure S5: Thy14 internucleotide NOE distributions in replicate 2. Acceptable distance ranges (shaded green) were calculated from the raw NOE data obtained from the RCSB. Graphs are labeled with residue1.atom1.residue2.atom2 in the top right of each graph.

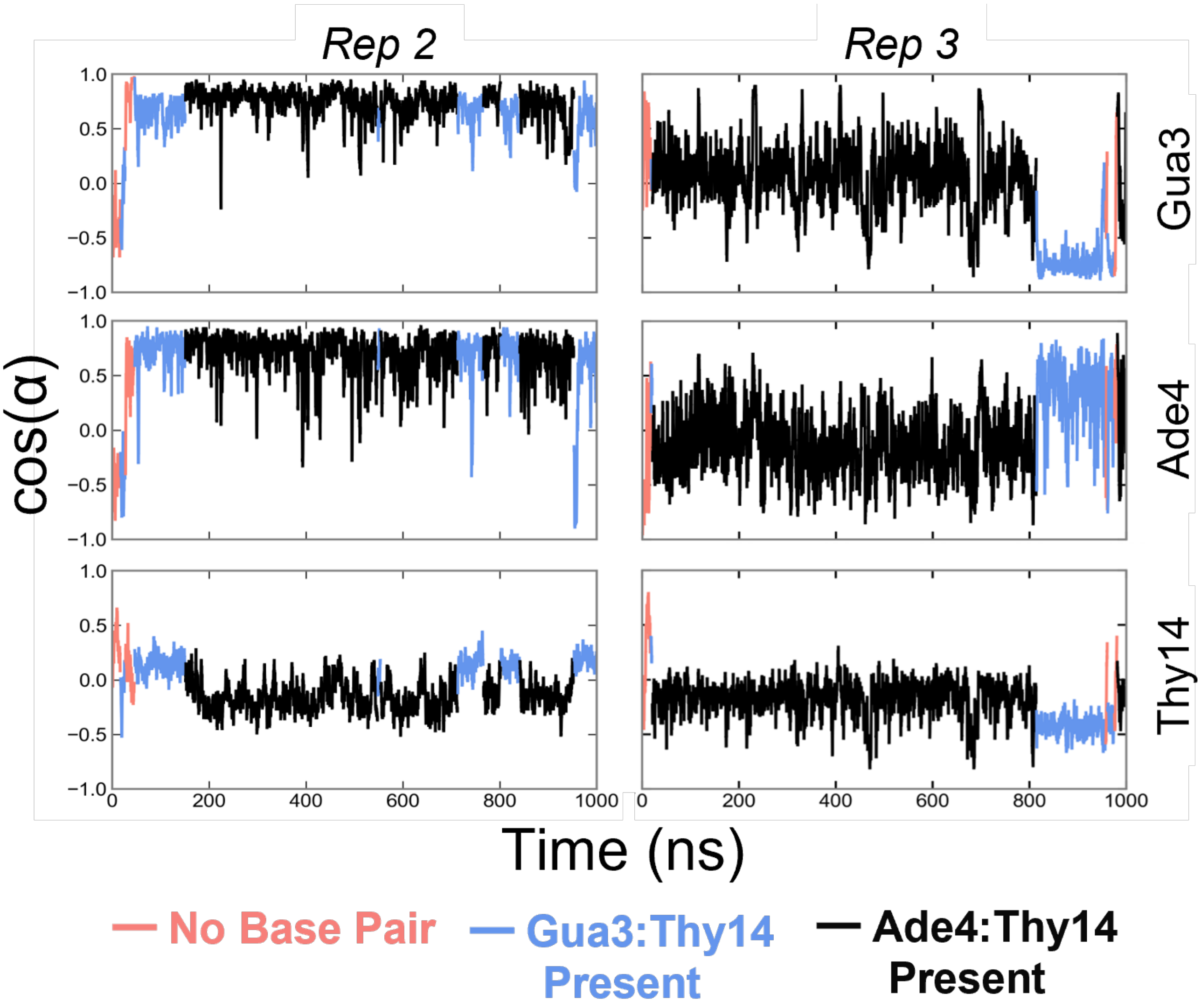


Figure S6: Electric field and dipole moment vector alignment without solvent contributions. Graphs represent the alignment as a cosine function of the angle during no base pair (pink), Gua3:Thy14 (blue), and Ade4:Thy14 (blacks) states.

| Angle between $\vec{E}$ and $\vec{\mu}$: α (º) | | | |
|---|---|---|---|
| | | *Rep 2* | *Rep 3* |
| **Gua3** | **No Base Pair** | 121 ± 30 | 125 ± 30 |
| | **Ade4:Thy14** | 94 ± 30 | 118 ± 20 |
| | **Gua3:Thy14** | 102 ± 30 | 143 ± 10 |
| **Ade4** | **No Base Pair** | 105 ± 40 | 108 ± 40 |
| | **Ade4:Thy14** | 86 ± 40 | 85 ± 30 |
| | **Gua3:Thy14** | 84 ± 40 | 77 ± 40 |
| **Thy14** | **No Base Pair** | 116 ± 20 | 113 ± 20 |
| | **Ade4:Thy14** | 113 ± 20 | 109 ± 20 |
| | **Gua3:Thy14** | 93 ± 20 | 111 ± 20 |

Figure S7: Average and standard deviation of the alignment between the electric field and the base dipole moment vectors for Gua3, Ade4, and Thy14 during all hydrogen bonding patterns: No base pair (pink), Ade4:Thy14 (grey), and Gua3:Thy14 (blue).

| Base Dipole Moment: \|μ\| (Debye) | | | |
|---|---|---|---|
| | | *Rep 2* | *Rep 3* |
| **Gua3** | **No Base Pair** | 8.35 ± 0.8 | 8.43 ± 0.5 |
| | **Ade4:Thy14** | 7.80 ± 0.6 | 8.03 ± 0.6 |
| | **Gua3:Thy14** | 8.05 ± 0.6 | 8.37 ± 0.6 |
| **Ade4** | **No Base Pair** | 3.74 ± 0.9 | 3.48 ± 0.8 |
| | **Ade4:Thy14** | 3.39 ± 0.7 | 3.41 ± 0.7 |
| | **Gua3:Thy14** | 3.51 ± 0.7 | 3.67 ± 0.8 |
| **Thy14** | **No Base Pair** | 6.59 ± 0.7 | 6.76 ± 0.6 |
| | **Ade4:Thy14** | 6.54 ± 0.5 | 6.44 ± 0.6 |
| | **Gua3:Thy14** | 6.62 ± 0.5 | 7.10 ± 0.6 |

Figure S8: Average and standard deviation of base dipole moments during all hydrogen bonding patterns: No base pair (pink), Ade4:Thy14 (grey), and Gua3:Thy14 (blue).

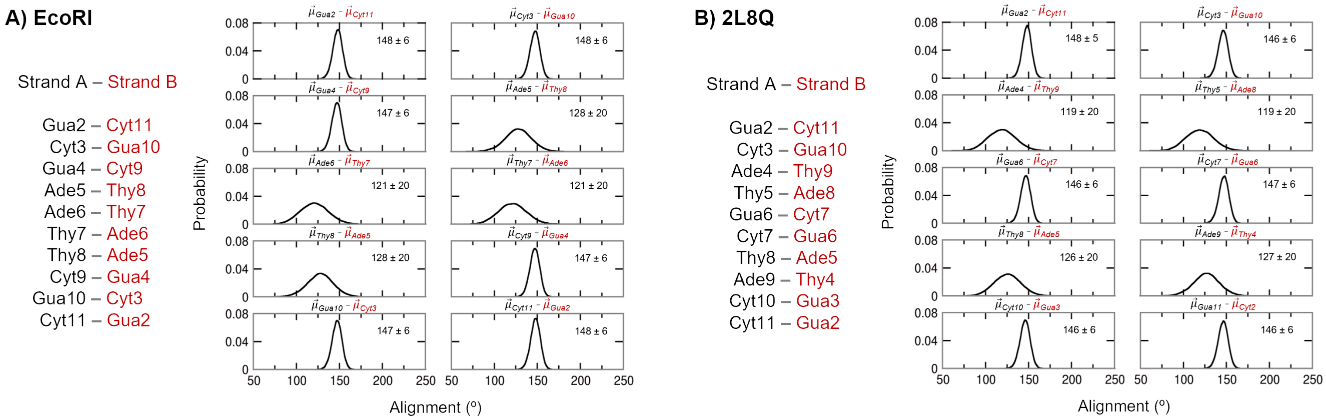


Figure S9: Model duplex base pair dipole moment alignments for MD simulations previously performed with the Drude-2017 FF of (A) EcoRI from PDB 1BNA and (B) the duplex from PDB 2L8Q.

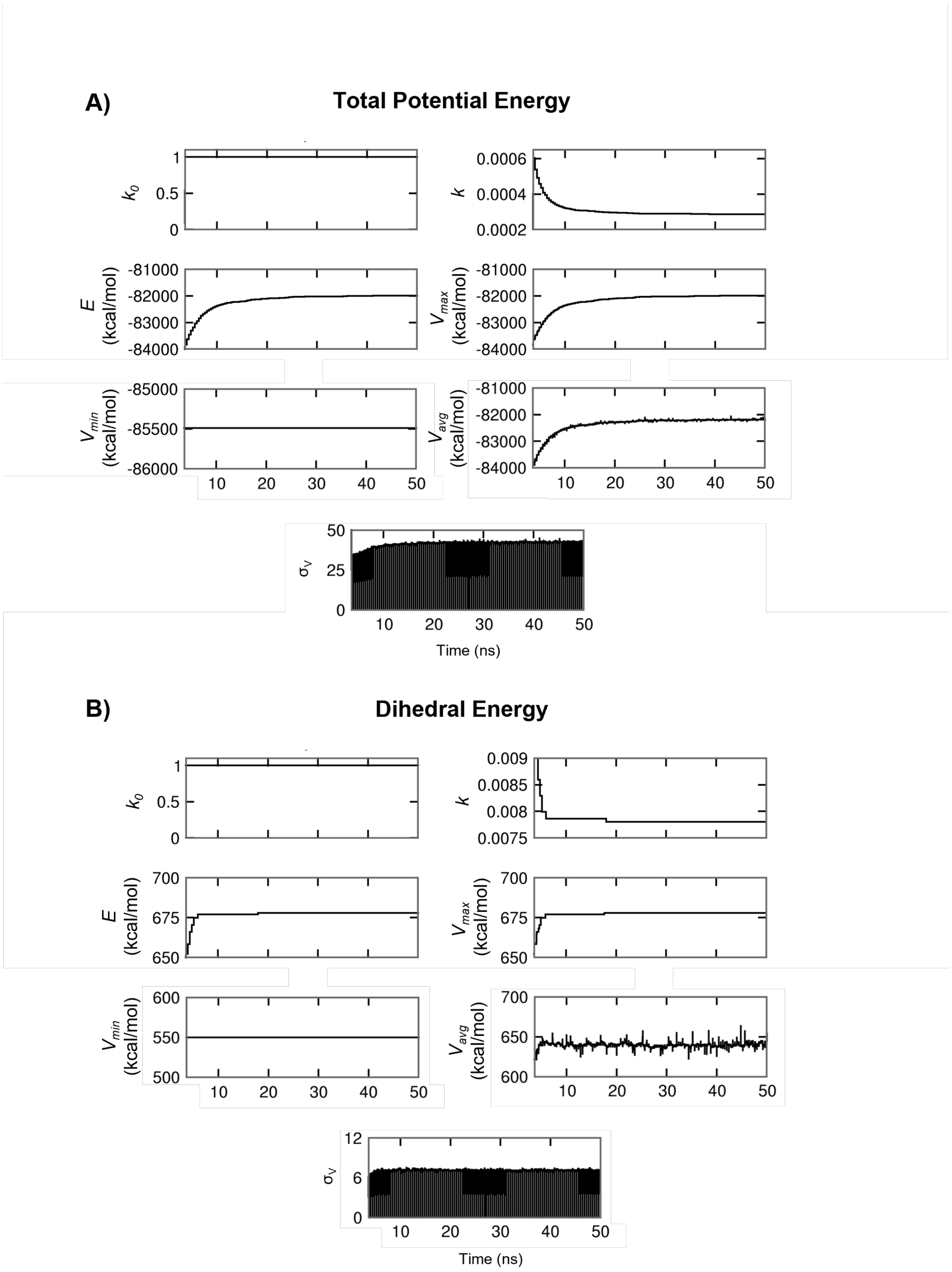


Figure S10: GaMD statistical information from equilibration. Both (A) total potential energy and (B) dihedral potential energy were evaluated for convergence.

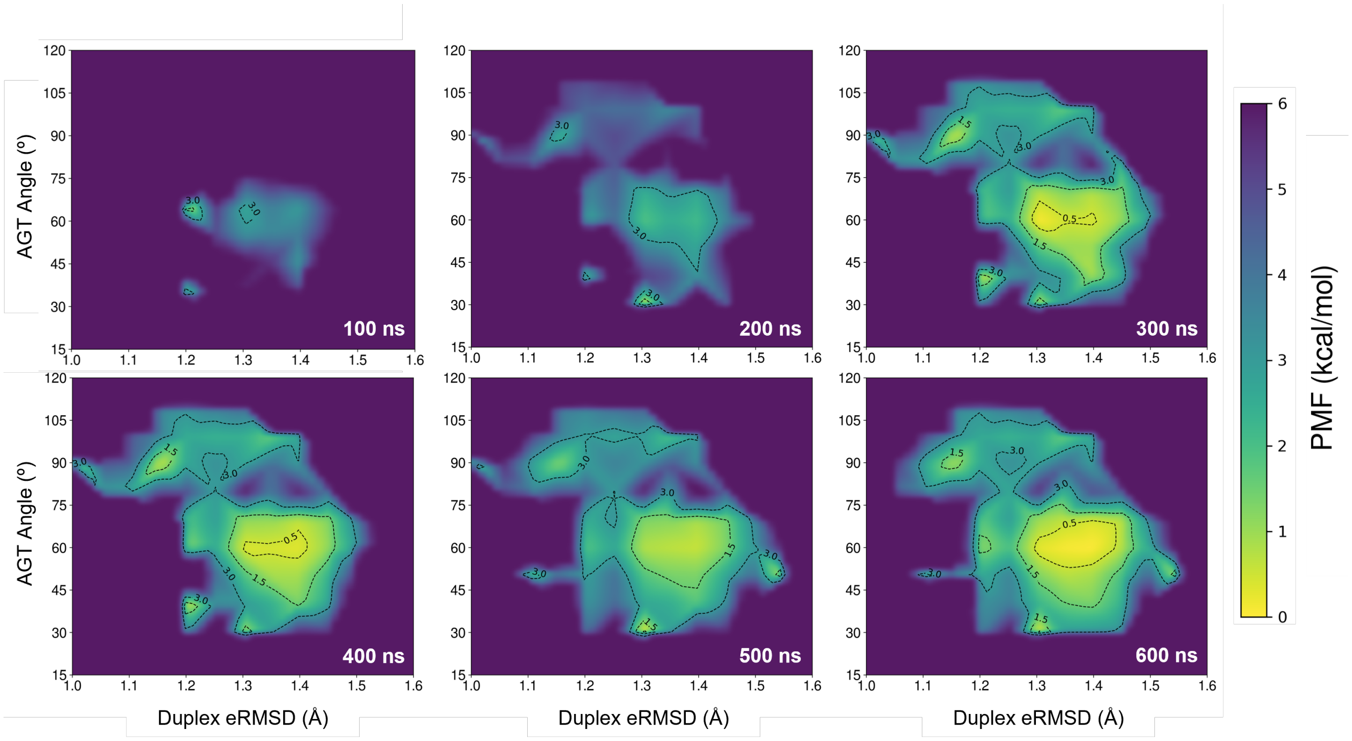


Figure S11: GaMD free energy surfaces over time. Data were pooled and reweighted every 100 ns using cumulant expansion to the second order.
